# Strengthening conservation law enforcement: incorporating harm, remediation and plural nature value criteria into an individual-based assessment of biodiversity offence gravity

**DOI:** 10.1101/2025.01.16.633328

**Authors:** Dominique Ghijselinck, Olivier Honnay, Erik Matthysen

## Abstract

Biodiversity loss is a pressing global issue, necessitating effective conservation laws and compelling enforcement mechanisms. Yet, implementation issues persist. This study contributes to the development of an individual-based decision tool designed to help judicial authorities accurately assess the gravity of biodiversity offences and the harm inflicted on individual organisms or a specific habitat. Through an extensive survey of expert opinions from 105 conservation biologists, we investigate what harm, remediation, and plural nature value criteria are most critical for the individual-based assessment of the gravity of biodiversity offences. We also identify potential pitfalls involved in the design of a decision tool. Experts identified species-specific traits such as specialization, generation length, and dispersal ability, as well as habitat characteristics like connectivity and natural integrity, as crucial harm criteria. A majority of experts also stressed the importance of remediation criteria that reflect the potential of harm reversal. Nevertheless, key challenges emerged, including the limitations of using single criteria, data insufficiency or uncertainty, and the complexity of combining and weighting these criteria into an overarching gravity score. Experts emphasized the need for a well-defined, distinct, and transparent set of criteria, suggesting that a blend of general and specific criteria might offer a pragmatic solution. Finally, our analysis also addressed value-laden implications, revealing that a plurality of underlying reasons for valuing biodiversity profoundly shapes how biodiversity offences are perceived.

## 1. Introduction

Biodiversity is declining globally at rates unprecedented in human history, with species extinctions continuing to accelerate (IPBES, 2019; De Witt et al., 2019). Conservation laws can be very effective means to mitigate biodiversity loss (Chapron et al., 2017; Phelps et al., 2021). For example, global and regional instruments, such as the European Birds (Directive 1979/409) and Habitats Directives (Directive, 1992/43) grant protection to a considerable number of taxa (Hunter et al., 2021) or, as in the case of the Environmental Liability Directive, define legally recognized harm caused to them (Directive, 2004/35). To attain their conservation objectives, these instruments must be duly transposed into species-oriented plans and strategies within domestic or regional legislative systems aimed to mobilize, guide and coordinate conservation efforts on a national level (Sazatornil et al., 2019). A key element for success is a strong enforcement with effective prosecution, conviction, and sanctioning of biodiversity misdemeanours, malpractices and felonies, with corresponding levels of legal repercussions and remediation or compensation (Bennett, 2011; Akella & Cannon, 2017).

Three important implementation issues with regard to conservation laws, hitherto largely unaddressed by the scientific community, relate to the assessment of the ‘gravity’ of biodiversity offences. With gravity, we here refer to the *degree* of different kinds of harm that biodiversity offences cause. Firstly, despite being deemed essential to an effective implementation of conservation laws, there currently exists no comprehensive individual-based decision tool to help the judiciary shape and implement prosecution and sentencing policies, or the sanctioning of biodiversity offences, in a proportional and dissuasive way (Billiet et al., 2018; Musing et al., 2018; Boogers et al., 2020). With ‘individual-based’, we refer to a particular methodological perspective that allows for the qualitative and quantitative assessment of the gravity of biodiversity offences down to the level of the killing, capture and possession of individual organisms, and damage to, or destruction of, individual habitat patches. Secondly, among those studies that assess the value of biodiversity more broadly, many use economic valuation frameworks to estimate its relative value based on market data, usually using a willingness-to-pay approach (Carson, 2012; Environmental Valuation Reference Inventory, 2024). However, a market value may be a poor reflection of the biological or intrinsic value of biodiversity (Soulé, 2014; Small et al., 2017; Washington et al., 2017; Vaissière & Meinard, 2021; Muradian & Gómez-Baggethun, 2021). More importantly, market-based approaches do not provide clear guidelines for how lawsuits could result in actions that meaningfully remedy harm to biodiversity and bring it back to its condition as closely as though the offence had not occurred (Phelps et al., 2021). This also aligns with the notion of “restorative justice” presented by Phelps et al. (2021) and Hutchinson et al. (2023). This concept describes how a lawsuit could ask a defendant to undertake actions to make amends for the harm caused to biodiversity, which also include more social-ecological impacts of biodiversity loss (Chan et al., 2016, IPBES, 2022). Finally, and more generally, several scholars have called for an interdisciplinary collaboration integrating insights from conservation biology, legal sciences, and ethics to enhance conservation law enforcement (Trouwborst et al., 2017; Chapron et al., 2017 ; Phelps et al., 2021 and Mateo-Tomás et al., 2022)

As such, there is a need for a decision tool aimed at the individual-based assessment of the gravity of biodiversity offences that comprehensively integrates gravity criteria into a concrete quantitative metric (Directive 2004/35; Directive 2008/99; Chapron et al., 2017; Naves et al., 2020; Phelps et al., 2021). This would allow a judge or administrative authorities to better substantiate sanctioning and remedial decisions, making these more effective in promoting biodiversity conservation (St John et al., 2012; Chapron et al., 2017; Billiet et al., 2020). We refer to a quantitative metric of gravity as a ‘gravity score’. For example, a higher gravity score, may then be linked to higher penalties or fines, confiscations, and non-custodial approaches such as community service, re-education, remediation measures, or financial compensation. Conceptually we distinguish between three different types of criteria that may contribute to such a gravity score. Harm criteria account for biological and ecological characteristics that could enable a more nuanced understanding of the degree of harm to the viability of populations and/or functioning of habitats. Remediation criteria account for the risks associated with remediation (reversibility of harm) and compensation (irreversibility of harm). Adding a plural nature value angle to the conceptualisation of gravity allows for the integration of diverse value perspectives into our analysis. Such value pluralism, or criteria derived from it, not only refers to nature’s contributions to human wellbeing in a material instrumental sense but also comprise the cultural, historical, recreational or aesthetic importance of natural places to people (Cervinka et al., 2012; Chan et al., 2016; Jimenez et al., 2021) or nature’s intrinsic value (Washington et al., 2017; Ghijselinck, 2023).

The general objective of this study is to contribute to the development of an individual-based decision tool to help judicial authorities accurately assess the gravity of biodiversity offences. Therefore, we used a survey of expert opinions of 105 European conservation biologists to: (i) investigate what harm, remediation and plural nature value criteria they consider most important for the individual-based assessment of the gravity of biodiversity offences; and (ii) identify potential pitfalls involved in the design of a decision tool that combines and integrates these criteria in a score reflective of the gravity of a given biodiversity offence.

## 2. Methods

### 2.1. Gravity criteria

To identify potential harm and remediation criteria, we took our cues from Environmental Impact Assessment literature. We specifically focused on literature calling for more comprehensive biodiversity metrics which account for habitat-level and population dynamic processes driving species occurrence and abundance, and remediation of biodiversity loss, into decision-making and policy (Quétier & Lavorel, 2011; Bigard et al., 2017; Marshall et al., 2020; Kujala et al., 2015; Maron et al., 2016; zu Ermgassen et al., 2019a; Marshall et al.,2020; Marshall et al., 2022; Fraixedas et al., 2022).

In our analysis of harm criteria, we distinguished between harming individual organisms and individual habitats. Harm criteria for habitats relate to the distinctiveness, quality or connectivity of the habitat patches concerned. Harm criteria for individuals encompass the conservation status of species, their role within an ecosystem, and concepts from population dynamics and life history. Note that the ‘individual-based assessment’ can also be supported by species-specific gravity criteria alongside individual- or context-specific factors, which respectively say something about characteristics specific to the individual entity involved in offences (e.g., age, reproductive value) and the location (e.g., in a protected area) or timing (e.g., during the breeding season) of the biodiversity offence. ‘Individual-based’ also needs to be distinguished from ‘individual-directed’, in that it not necessarily means that the loss of the individual organism as such, and correspondingly its intrinsic value as subject-of-a-life or end in itself in a moral sense is deemed important (Himes & Muraca, 2018). An individual-based assessment of the gravity of biodiversity offences can therefore also be ‘species-directed’ conveying the importance of the value of biodiversity on the level of populations or species.

We included remediation criteria because remediation is increasingly part of legal requirements or policy obligations (Phelps et al., 2021; Droste et al., 2022). As Phelps et al. (2021) explain, remedy-focused liability litigation is an important conservation strategy that creates novel approaches for justice, while compensation can deliver financial support for conservation. Such reasoning aligns with the ‘polluter pays’ principle, where the person who causes a loss must, in addition to punishment, bear the burdens of remediating, or compensating, the party that suffers it, which is also the declared objective of the EU Environmental Liability Directive (Directive 2004/35). To establish remediation criteria we employ the ecological, geographical, and temporal flexibility of remedial actions which respectively pertain to how, where and when to remedy harmed biodiversity. These flexibilities are an adaptation of the three types of remedial measures as presented in the EU Environmental Liability Directive (Directive 2004/35). In terms of the ecological flexibility of remedial action, primary or in-kind remediation at the same site is preferable (Directive 2004/35). ‘In-kind’ remediation aims to bring the ecosystem’s structure back to its condition as closely as though the offence had not occurred. As flexibility increases the communities or species targeted by remediation can be increasingly different from those impacted by the offence (Bull et al., 2015). Geographical flexibility pertains to how, if primary remediation cannot fully restore the damaged site to its baseline condition, remediation measures must be adopted at a different and more distant location (Directive 2004/35; zu Ermgassen et al., 2019b). Temporal flexibility pertains to interim losses and how harmed biodiversity is not able to perform ecological functions or provide services until remedial measures have achieved their full effect (Directive 2004/35; zu Ermgassen et al., 2019b).

In addition to harm and remediation criteria, we incorporate a plural nature value perspective to expand the discussion on the gravity of biodiversity offences. This approach recognizes that diverse judgments about nature’s importance shape different views on human-nature relations, influencing conservation framings and decisions (Mace, 2014; Pascual et al., 2022; Ghijselinck et al., 2023). We base this perspective on the IPBES (2022) typology of nature’s values, further developed by Anderson et al. (2022), which classifies nature’s value into instrumental (nature as a means to an end, often associated with ecosystem services), relational (meaningful reciprocal human-nature relationships), and intrinsic (nature deserving moral consideration in itself).

### 2.2. Survey of European conservation professionals

#### 2.2.1. Participant identification

We focused our study on European countries and contacted conservation biologists based in Europe, to encompass diverse views within the conservation researchers’ community there. We considered experts as people with a doctoral degree in this field and actively participating in research on biodiversity conservation. To find contact addresses, we consulted the *Science Direct* and *Web of Science* search engines to track down post-2019 publications on biodiversity and conservation related topics, and identify the lead authors of these publications. As search terms we used ‘conservation’ and ‘fauna’, ‘flora’, ‘biology’, ‘ecology’, ‘biodiversity’, and looked for their presence in titles and abstracts.

#### 2.2.2. Design of the questionnaire

Following Ritchie and Ormston (2014), we used a quantitative and qualitative in tandem approach to elicit distinctive types of data. We employed closed-ended questions, asking respondents to indicate their personal levels of agreement on a 5-point Likert scale. A few questions included a ‘no opinion’ option. We also attempted to get additional insights into the respondents’ opinions and asked to elaborate on answers given to the closed-ended questions. For some questions we also introduced sample topics with a short explanatory text or case study to ease interpretation and understanding of the statement. Table 1 presents the surveyed harm and remediation criteria we used to structure the questionnaire, and shows the rationale providing the grounds for why a given gravity criterion can be used to increase (or lower) the gravity score of a biodiversity offence. We used the same format to analyse the gravity of biodiversity offences from a plural nature value perspective, eliciting respondents’ opinions on the importance of landscape elements that are locally or regionally recognised as part of natural or cultural heritage, or have aesthetically appealing properties. A few questions enquired about whether, in the respondents’ opinion, a gravity score for a given offence should go beyond the integration of biodiversity values that affect human wellbeing (instrumental value) to also include relational and intrinsic notions. We also asked about the need for a balance between the use of specific criteria versus general ones in assessing the gravity of biodiversity offences. A copy of the full survey can be found in the Supporting Information.

**Table 1:**
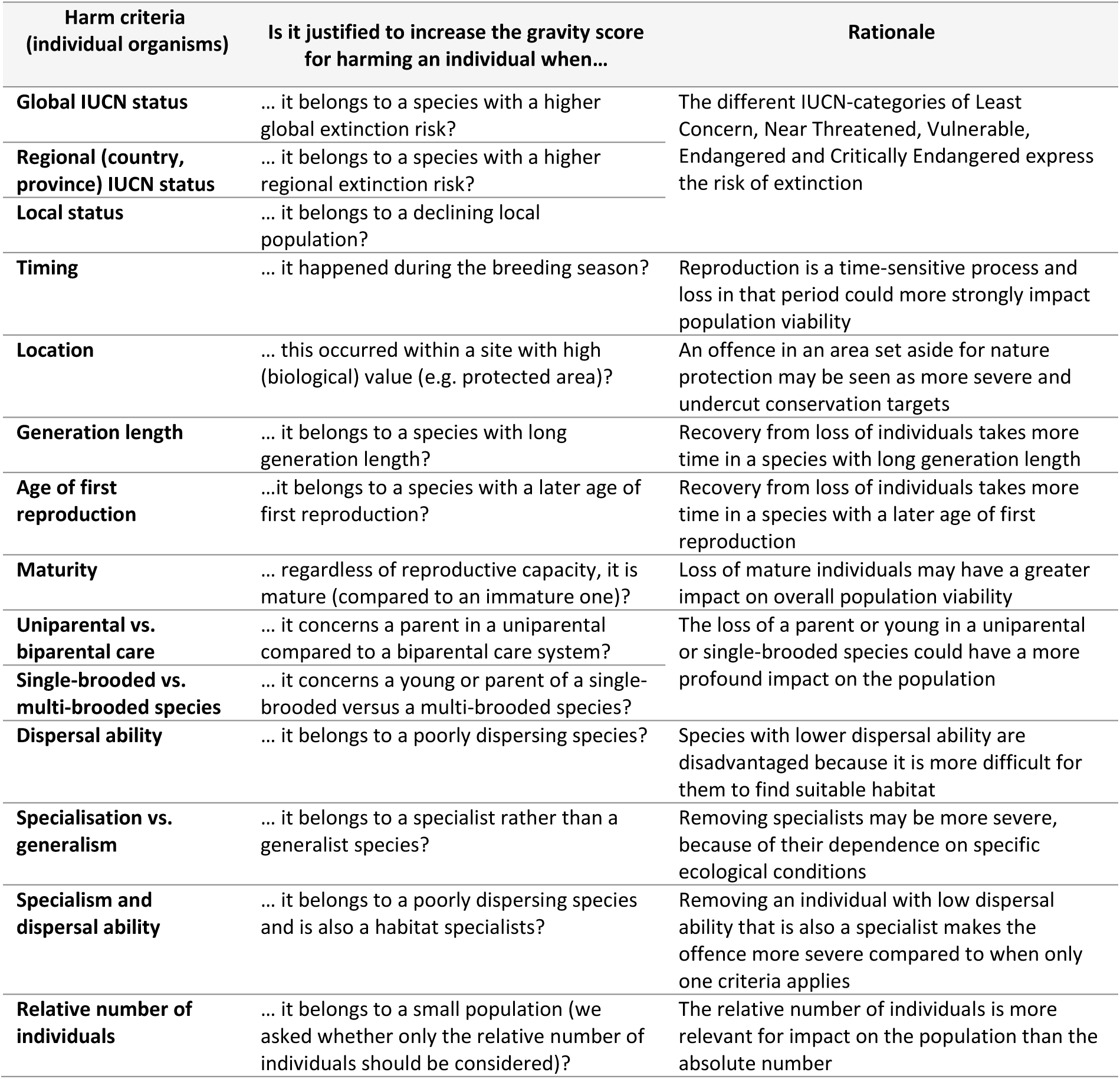
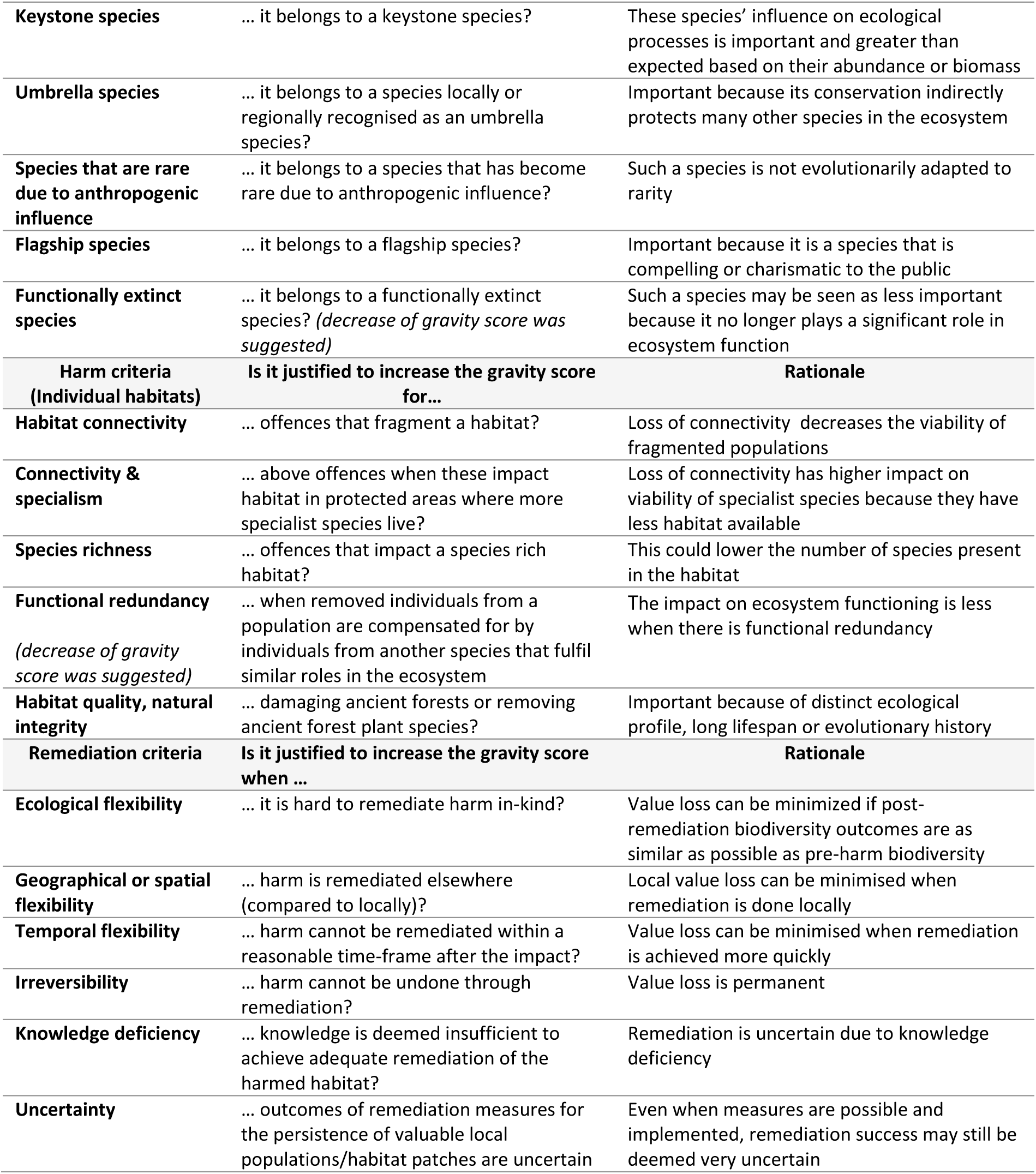
Harm and remediation criteria addressed in the survey. The middle column presents a simplified representation of the survey questions, while the rightmost column shows the rationale for using a given gravity criterion.

#### 2.2.3. Procedure and ethics

We collected data between mid-November 2023 and February 2024, and administered the survey online by using the Qualtrics Software Survey Platform (Qualtrics, n.d.). We invited 300 potential respondents to complete the survey by sending them an email with a weblink to the survey, of whom 105 responded. We sent two reminders at most to persons who did not reply. Before launching, three persons actively involved in biological conservation research pretested the survey. This further led to suggestions and modification of the survey and filtered out questions or statements that were considered too abstract.

We screened and analysed survey data quantitatively using the Qualtrics Results tab. In the few cases where the answers given suggested misinterpretation or contradiction we sent a follow-up question to ask for clarification. We utilised descriptive statistics, such as percentages, means and standard deviations to summarize the Likert-data collected. Because the sample of conservation biologists participating in the survey was small and purposive we mapped the range and diversity of answers as well as providing quantitative statistics.

## 3. Results

The respondents showed considerable agreement on most of the harm criteria relating to the gravity of biodiversity offences to individual organisms (Figure 1). We found most agreement (SD<1) on the relevance of regional IUCN and local status of species, and harmed individuals from specialist and keystone species, or species with a long generation length, older age at first reproduction and low dispersal ability. Other responses are more widely spread, suggesting varied opinions. This is the case for some concepts that primarily pertain to life history strategies, the timing of the offence, conservation surrogate species such as flagship and umbrella species, and, remarkably, global IUCN Red List status. As for harm criteria for habitats, responses indicated a high agreement for the relevance of criteria related to ecological connectivity and natural integrity and more spread for species richness or functional diversity (Figure 2). For all surveyed remediation criteria we found a high agreement (SD< 1), whereby agreement was highest for an increase of the gravity score in cases of harm that is irreversible, and when remediation is not in-kind or takes a lot of time.

**Figure 1:**
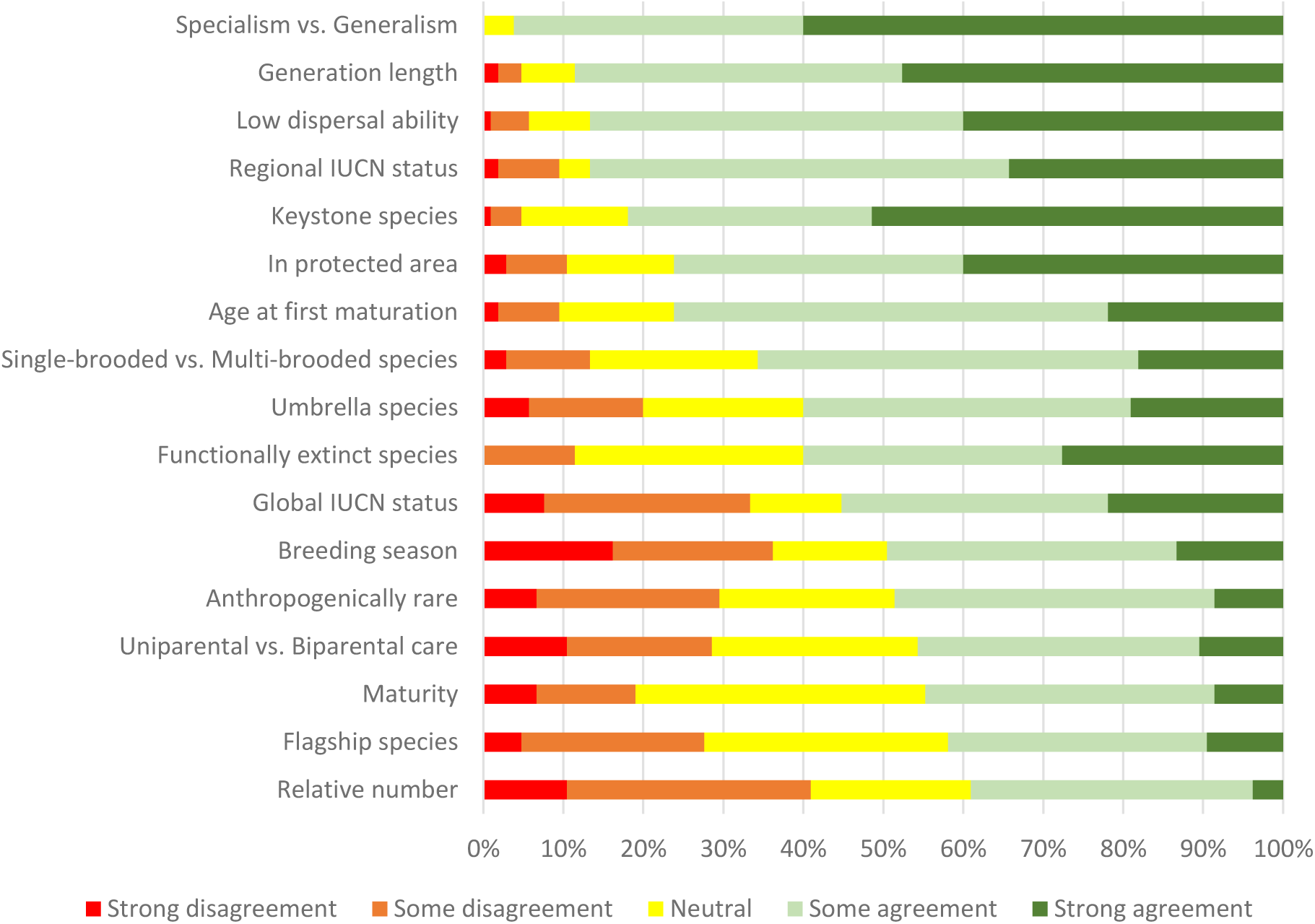
Stacked bar chart showing harm criteria for individual animals or plants ranked by agreement. Agreement is linked to an increase of the gravity score, except for ‘functionally extinct species’, where scores were reverse-coded.

**Figure 2:**
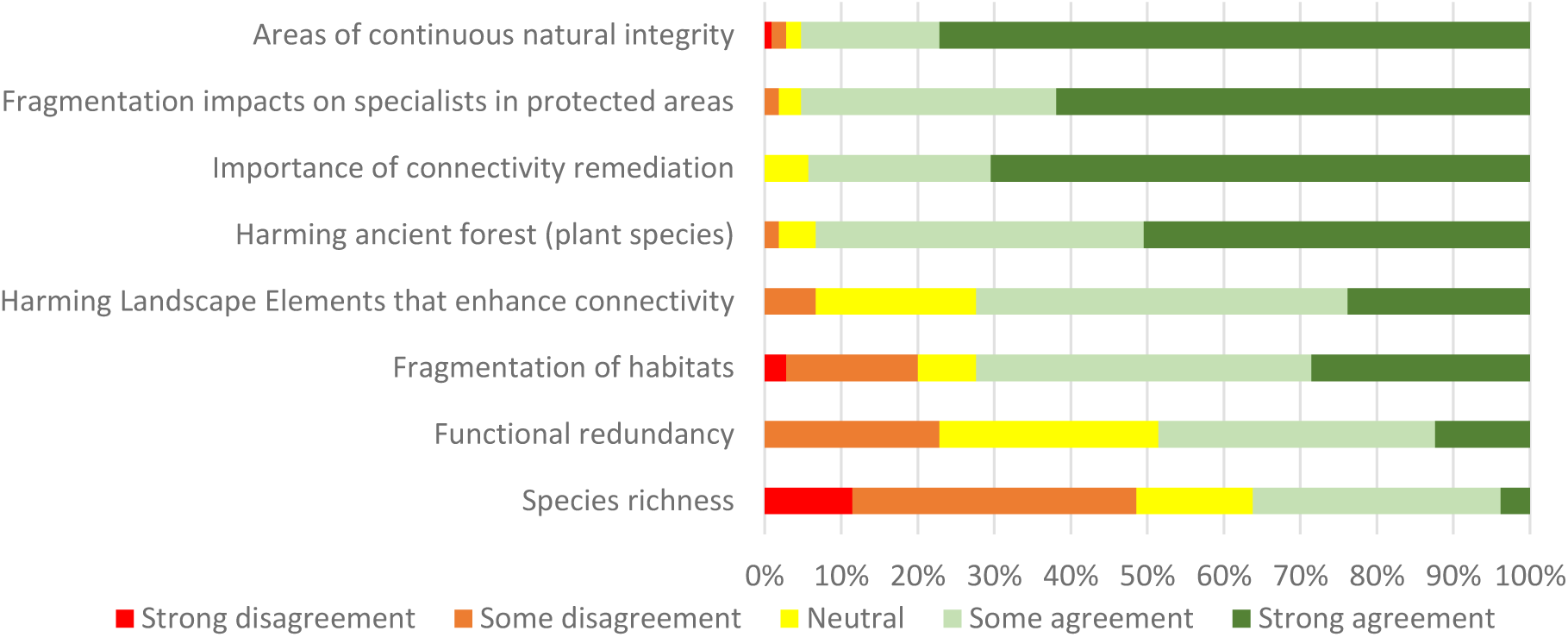
Stacked bar chart showing suggested harm criteria (habitats) ranked by agreement. Agreement is linked to an increase of the gravity score, except for ‘functional redundancy’, where scores were reverse-coded.

In the following text, unless otherwise mentioned, ‘agreement’ or ‘support’ refers to the combination of ‘some agreement’ and ‘strong agreement’.

### 3.1. Survey data: harm criteria for individual organisms

#### Conservation status

About half of the respondents (55%, mean =3.36, SD=1.28) supported the use of global IUCN Red List status as criterion, while this was substantially higher for Regional status (87%, mean=4.10, SD=0.92). Most respondents found that when the individual harmed is from a species with poor local status or, when not available, declining local population trends, this should increase the gravity to some extent (35%) or to a large extent (50%).

Many respondents pointed out the global status may not be reflective of the regional or local population statuses. Several respondent mentioned that the latter are more important in assessing the gravity of an offence or even that harm to individuals within a small population is more damaging, regardless of the conservation status. Nonetheless, even when the individual harmed locally belongs to a species of which the local conservation status is favourable, 81% of respondents found that the gravity score should be increased at least to some extent when the species is threatened in other parts of its range. Importantly, some respondents pointed to the lack of data, especially regarding the conservation status of less well-known taxa (e.g. insects). In this case, some respondents added local expert judgement and verification may be warranted.

#### Timing and location of impact

Only half of the respondents (49.5%, mean=3.10, SD=1.32) agreed that gravity scores should be increased for the killing of a mature animal during the breeding season compared to other times of the year, while 16% strongly disagreed and 20% somewhat disagreed. Remarkably, the only argument offered by respondents who disagreed, was the complexity of applying this criterium. A majority of 76% (mean=4.03, SD=1.05) agreed to increase the gravity score for offences occurring within a site with high (biological) value (nature reserve, Natura 2000, National Park), while a minority (10%) disagreed.

#### Generation length, age at first maturation

Over 88% (mean=4.3, SD=0.86) of respondents agreed to use generation length to assess gravity when the individual harmed belongs to a local population which is declining. According to 76% (mean=3.87, SD=0.91) the age at first maturation can be used as an alternative for generation length.

However, among those who agreed, some expressed concerns about the use of generation length as a harm criterion, and pointed out that it is already used as such in the IUCN Red List. Indeed, population reductions are scaled with generation length by the IUCN because species with longer generation length recover more slowly from declines, whereas rates of human-induced declines are not limited by biological constraints (IUCN, 2022).

#### Maturity

Less than half of respondents (45%, mean=3.28, SD=1,01) agree that, regardless of reproductive status, a gravity score should be higher for a mature compared to an immature individual of the same species. Some respondents underscored how, in addition to their breeding capacity, mature individuals often have more experience in finding food, avoiding predators, and navigating their environment, while immature individuals are still in the process of developing and honing their survival skills. Nevertheless, some respondents stated this was too complicated to use as a harm criterion.

Several respondents argued that reproductive value (i.e. the expected future breeding output of an individual) could also be used as an individual criterion, because the loss of an animal with a high residual reproductive value could potentially be more detrimental to the population’s health and viability. However, at the same time it was pointed out that this would require detailed information about age-specific reproductive output, survival probabilities, and the future reproductive potential of both the individual and its offspring. Therefore, it was argued such information need is probably too comprehensive to be used in a legal context or else a proxy could be used.

#### Uniparental versus biparental care

Using birds as an example, respondents were asked whether the gravity score for harm to a parent bird should be higher in case of uniparental care than in the case of biparental care. Less than half of respondents agreed to this (46%, mean=3.17, SD=1.16). Some respondents again remarked that this was too complicated as harm criterion, because, in summary, one should then consider the unique characteristics of each species and the potential consequences of losing a parent in different caregiving systems. Also, according to one respondent:

> Biologically it is reasonable, in the case of biparental care, the nest still would have some chance of success. However, in practice, it would then start depending on whether the nest loss was discovered or not, which may not be an optimal sanctioning rationale.

#### Single-brooded versus multi-brooded species

Nearly two-third of respondents (66%, mean=3.68, SD=0.98) agree that the killing of young or a parent should lead to a higher gravity score in a single-brooded versus a multi-brooded species. Here, one respondent commented they:

> strongly agree for a fledgling because there would be no more fledglings produced in the same breeding season in case of a single-brooded species, but somewhat disagree for a parent, because it would probably have consequences across breeding seasons irrespective of single or multiple brooding.

#### Dispersal ability and specialist species

Most respondents (87%, mean=4.20, SD=0.87) agreed it is justified to increase the gravity score for harm to an individual of a poorly dispersing species regardless of whether its habitat is currently fragmented or not. However, many of those who agreed remarked it may be difficult to properly identify a species’ dispersal ability.

Most respondents (96%, mean=4.56, SD=0.57) agreed that, in general, harm done to habitat specialists should be given more weight compared to generalists. Respondents pointed out that, as they have more special requirements, specialists are more at risk than generalists, and are more vulnerable to habitat isolation. Likewise, there was near consensus (91%, mean=4.36, SD=0.66) that a limited dispersal capacity of harmed habitat specialists should be seen as an additional gravity factor (on top of specialism). Some respondents commented that specialisation, in the case of a high local abundance, need not lead to an increase of the gravity score.

#### Number of individuals harmed

There was strong divergence of opinions (39% agreed; 41% disagreed) as to whether only the relative number of harmed individuals should be taken into account instead of the absolute number. Among those who disagreed, some remarked they would agree when the number of individuals harmed would negatively impact the whole population, but that it may not be easy to discern when this is the case. In general, many respondents pointed out that the loss of an individual in terms of population viability does not matter if the population and habitat of the species concerned are not threatened and in good condition.

#### Ecological role

Most respondents (82%, mean=4.28, SD=0.90) agreed that harm to an individual belonging to a keystone species should lead to a higher gravity score compared to other species. However, over one third (34%, mean=2.86, SD=1.15) agreed defining keystone species may be difficult and therefore this criterion should be used with caution. Only 60% (mean=3.53, SD=1.12) of respondents agreed on a higher gravity score for umbrella species.

Respondents were divided on whether harm done to an individual from a species that has become rare due to anthropogenic influence is worse compared to a naturally rare species, with 40% agreeing somewhat, 8.5% strongly agreeing and 22% remaining neutral (mean=3.21, SD=1.09). One respondent remarked that:

> … the essence of this debate is that the conservation of both artificially and naturally rare species is crucial for maintaining biodiversity and that other criteria such as conservation status and local abundance may be better placed to distinguish between these two types of species.

We found both agreement (42%) and disagreement (27%) as to whether it is justified to increase the gravity score for a flagship species (mean=3.19, SD=1.04). This was also supported by diverging arguments. Some respondents remarked that the concept of flagship species works to engage with people but that it should by no means affect conservation decision-making and has poor correspondence with conservation status. Also, several times it was mentioned that by focusing on flagship species, species that are less appealing to a general audience but are more important for ecosystem functioning may be overlooked or not taken into account.

When the individual harmed belongs to a species which is functionally extinct only 11% (mean=2.24, SD=0.98) of respondents somewhat agreed this should lower the gravity of the offence.

### 3.2. Survey data: harm criteria habitats

#### Habitat connectivity

We presented respondents with a scenario in which protected dunescapes could be found interspersed by semi-natural or urbanised areas. The scenario assumes further fragmentation of these dunescapes could lead to detrimental impacts on biodiversity. Our questioning wanted to elicit the relevance of connectivity-related harm criteria in assessing the gravity of offences against habitats. Over 70% (72%, mean=3.78, SD=1.12) of respondents agreed that, regardless of the dispersal ability of the species present, it is justified to increase the gravity score for activities that fragment a habitat. Furthermore, nearly three quarters of respondents agreed (72%, mean=3.90, SD=0.84) that it is justified to increase the gravity score for damaging or destroying nature elements such as hedgerows, trees or shrubs that may facilitate dispersal between patches of habitat.

Moreover, 94% (mean=4.65, SD=0.59) agreed remediation measures should be imposed for offences that fragment protected habitat patches or increase isolation of protected areas, even when these happen in unprotected zones. There was a near consensus (95%, mean=4.55, SD=0.65) that possible impacts from such activities on habitat specialists in protected areas should be taken into account when assessing the gravity of biodiversity offences. This last finding suggests that the presence of specialist species in affected habitats can also aid in the assessment of biodiversity offences that impact habitat (connectivity). However, respondents pointed out that linking connectivity loss to specific impacts on specialist species may, in practice, not be straightforward, making it difficult to assess, let alone quantify, the gravity of offences.

#### Species richness, functional diversity

Almost half of respondents (48,5%, mean=2.80, SD=1.12) found species diversity on its own a poor indicator to assess harm to a given habitat patch. Many respondents remarked that species richness, defined as the number of species in an area or habitat patch, differs fundamentally between habitats and that changes in species richness may not be immediately clear. One respondent remarked:

> I think this is very context dependent. E.g. if we have naturally species-poor systems (many higher latitude or isolated island systems), changes in species richness are often likely to be hardly any.

Also, several respondents argued that establishing the presence of rare or specialist species and impacts on these species is more relevant and that neither the abundance nor the functional roles in a community are indicated through a total species count. One respondent suggested perhaps Shannon diversity could be used because not just the number of different species (species richness) but also their relative abundances in a community (evenness), is taken into account.

To the question whether the gravity score for removing individuals from a population should not be increased when these are compensated for by individuals from another species that fulfil similar roles in the ecosystem at hand nearly half of respondents disagreed (12% strongly, 36% somewhat, mean=2.62, SD=0.97). Finally, some respondents thought that the inclusion of metrics of functional diversity, even species richness, into a decision tool to assess the gravity of biodiversity offences would be too complicated.

#### Habitat quality, natural integrity

A majority of respondents (93%, mean=4.42, SD=0.67) agreed it is justified to increase the gravity score for damaging ancient forests or removing ancient forest plant species (50.5 % strongly). Also, over three quarters of respondents strongly agreed (77%, mean=4.69, SD=0.69) this should be the case for damaging an area or site with a long history of continuous natural integrity (e.g., old-growth forest, freely running river system), while 18% somewhat agreed. However, the arguments in support of this agreement varied. Respondents mentioned that such areas will take a higher cost for recovery, that the potential to reach a similar status will be reduced after impacts and even that their remediation or compensation would be impossible. These arguments are also reflected in the findings on remediation criteria in the next section. Other respondents pointed out that these areas are important for their “rarity”, or that they “deserve special attention”, or have “potentially higher scientific and historical value” or are “ecologically unique”. One illustrative quote here is:

> This type of areas are becoming rarer every day and provide unique ecological but also symbolic benefits. Thus the gravity of its loss should be more dramatic.

### 3.3. Remediation Criteria

The majority of respondents (88%, mean=4.38, SD=0.86) agreed that, when offences lead to loss of nature components and it is hard to remediate this loss in-kind, it is appropriate to increase the gravity score (see Figure 3). Almost all respondents disagreed (95%, mean=1.24, SD=0.72) with the statement that when an old forest patch is logged and replaced by planting a forest of the same size, the lost ecological value has been remediated. In fact, according to 93% of respondents (62%, strongly agreed; 31% somewhat agreed, mean=4.55, SD=0.62), the gravity score should increase the more time it takes to achieve an equivalent ecological state of the lost forest. A similar majority was found in the case of harm done to nature components in general, with 89.5% (mean=4.31, SD=0.87) of respondents finding it appropriate to increase the gravity score when impacts from illegal activities cannot be remediated within a reasonable timeframe.

**Figure 3:**
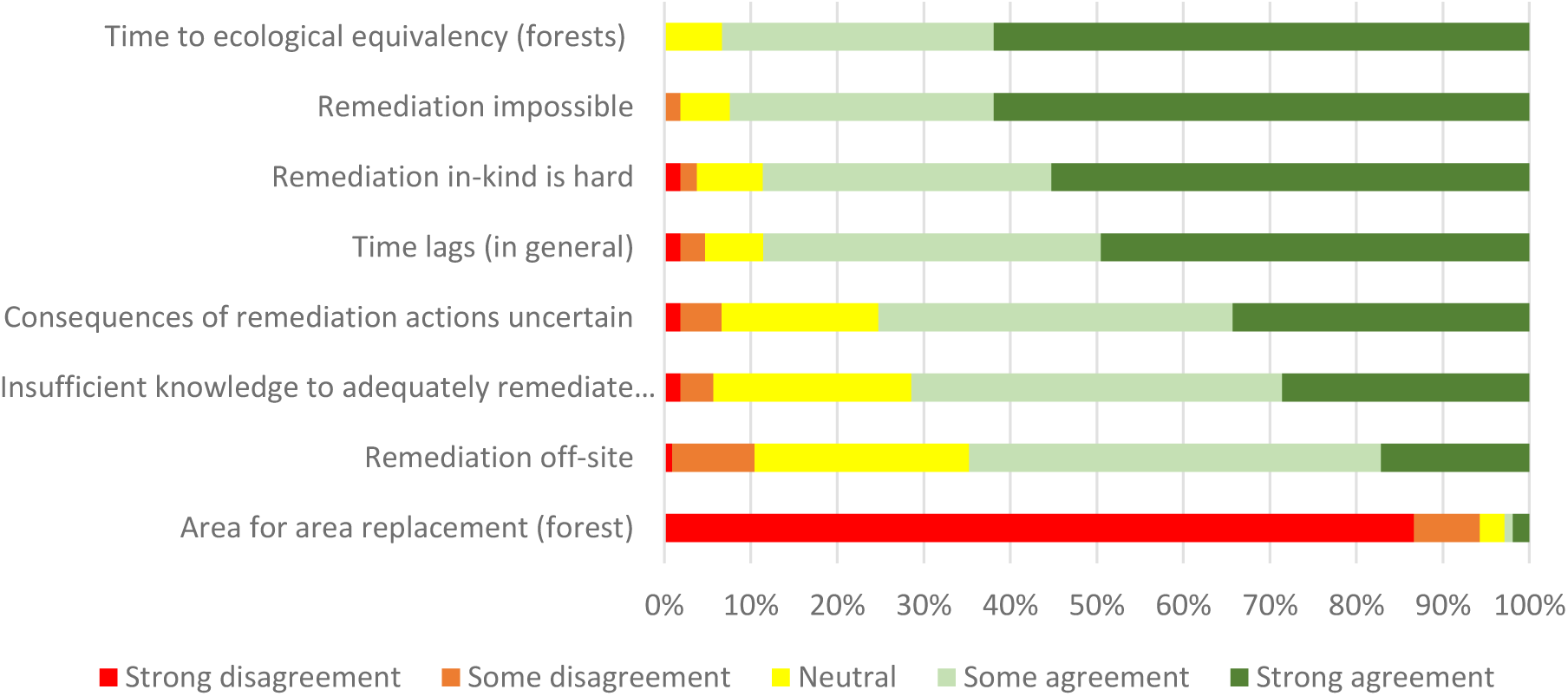
Stacked bar chart showing suggested remediation criteria ranked by agreement. Agreement is linked to an increase of the gravity score. Note that a majority of respondents considered area for area replacement of a lost forest inadequate as remediation measure.

Several respondents are of the opinion that shortcomings of, or risks associated with, pursuing remediation warrant complementary measures or compensation. When the consequences of remediation measures for the persistence of biologically valuable local populations or habitat patches is deemed uncertain it is appropriate to increase the gravity score beyond the efforts for remediation according to 75% (mean=4.01, SD=0.94) of respondents. In line with this, when knowledge of affected habitats is considered insufficient to achieve adequate remediation of habitat destruction, 71.5% (mean=3.91, SD=0.91) of respondents agreed this should increase the gravity score. Some also added that remediation measures can never fully return the community and functions to their baseline condition and that a higher gravity score should always imply complementary measures or compensation.

When asked whether increasing the gravity score is appropriate when remediation happens *elsewhere* (compared to locally), because local people lose the benefits this nearby natural zone provided, nearly two thirds of respondents agreed (17% strongly, 48% somewhat, mean=3.70, SD=0.89).

Most respondents agreed (92%, mean=4.52, SD=0.69) that when the removal of an unspecified native vegetation or killed individual is permanent - in that the harm is deemed irreversible and remediation in natura not possible - this should further increase the gravity score, beyond the one for remediation, and even when the offence happens in areas that have no protected status.

### 3.4. Plural nature value criteria

Just 21% (mean=2.59, SD=0.99) of respondents somewhat agreed that remediation of only instrumental values, presented as provisioning and regulating services, post-harm is sufficient (see Figure 4). Perhaps not surprisingly in light of this finding, respondents mentioned a variety of other than these instrumental reasons why they think the conservation of nature is valuable and why the loss of populations, species or habitats is undesirable. References to intrinsic value were made by over 40 people. The exact words used varied from simply “intrinsic value”, “inherent value” and “value on itself” or “non-anthropocentric value” to “out of respect”, “moral obligation”, “ethical reasons for not destroying the habitats and existence of other species” and “no right to destroy it [nature]”.

**Figure 4:**
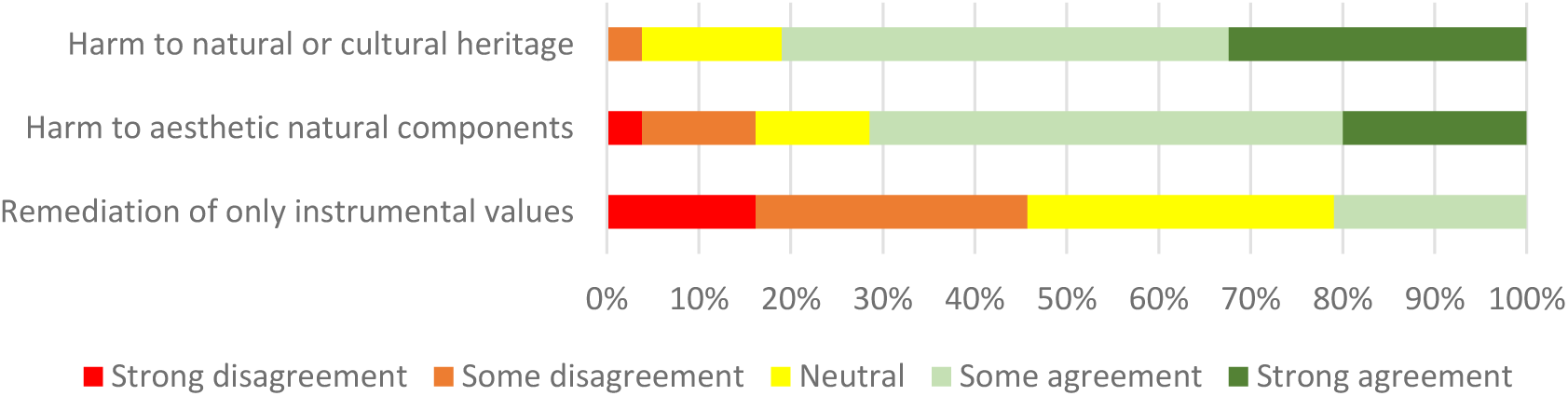
Stacked bar chart showing suggested values-based criteria ranked by agreement.

Respondents also mentioned recreational (spending time in nature, relaxation), aesthetic, educational (curiosity, learning), intergenerational and cultural identity values (how meaning and value is rooted in the land). Moreover, 81% (mean=4.10; SD=0.72) of respondents agreed that increasing the gravity score of an offence is justified when landscape elements are harmed that are locally or regionally recognised as part of natural or cultural heritage (such as hedgerows, shrubs and historical grasslands that have been part of the agricultural landscape for a very long time).

Furthermore, 71.5% (mean=3.71; SD=1.04) of respondents found an increase of the gravity score is justified when aesthetic properties of landscape elements, or feelings of reverence or awe, are involved (such as aesthetically appealing and very old, landmark trees). In general, these respondents remarked that ecological concerns, in terms of conservation targets, and intrinsic concerns should outweigh the consideration of social-ecological values in the assessment of biodiversity offences. In the words of one respondent:

> Social and human wellbeing aspects are most of the time the focus and most often result in the destruction of habitats. Humans are drawn to ameliorate their own well-being but often neglect the importance of ecological intactness.

## 4. Discussion and conclusion

We identified a set of gravity criteria that garnered majority, and in some cases, high agreement (over 85%) among the surveyed experts. These criteria can be used to support the individual-based assessment of the gravity of biodiversity offences. High agreement especially relates to criteria that are directly based on ecological processes or mediate population persistence - such as generation length, specialism, dispersal ability and connectivity metrics - and regional or local status. This, perhaps surprisingly, was not the case for global IUCN status, species richness or the timing of the offence. Similarly, there was strong consensus on the importance of remediation criteria, particularly those related to in-kind remediation and the time it takes to remediate harm. This underscores the need to use a diverse set of criteria to bolster an accurate and comprehensive assessment of biodiversity’s value and the gravity of harm inflicted. Despite the support we found for individual criteria, many respondents expressed reservations about the design of an individual-based decision tool that combines these distinct harm, remediation and plural nature value criteria into an overarching gravity score of a biodiversity offence, suggesting it may be feasible given the appropriate context and stipulations in which to use these criteria. Also, some respondents stated that more criteria do not necessarily mean more legal transparency, especially when data gathering is complicated or needs a lot of expertise.

We here revisit the concerns raised by the experts, implicitly or explicitly, regarding the use of gravity criteria and the design of a decision tool. We also offer recommendations to consider in decision tool design.

### 4.1. Relevance of gravity criteria: practical and pragmatic implications

Even when criteria were considered acceptable, respondents highlighted obstacles in collecting the ecological and other relevant data necessary to ensure the reliability and consistency of the decision tool. First of all, several respondents pointed to the lack or required comprehensiveness of data, as, for example, in the case of the conservation status for less well-known taxa, the determination of the reproductive value of an individual, the dispersal ability of a species, or a perceived keystone species’ actual impact. Another example here pertains to maturity, regardless of reproductive status. Respondents who supported the relevance of this criterion highlighted the specific role of the mature individual, noting that harming it could more significantly affect the overall stability of the population. Indeed, as is discussed by McDonald et al. (2020), many species exhibit complex social structures where mature individuals play crucial roles in maintaining social order, providing guidance, and contributing to the overall well-being of the group. Nonetheless, as respondents also admitted, this role may be hard to discern.

Moreover, respondents pointed out the application of criteria could be context-specific and emphasized that important qualifications or conditions must be considered when using criteria individually, as this may also impact the combination of criteria in an overall gravity score. This is the case with, for example, global conservation status, where respondents’ answers suggested a gravity score for harming an individual organism should at least somewhat increase when the species is threatened in other parts of its range, while on itself, this status was deemed insufficient in assessing the gravity of offences on a regional or local level by nearly half of the respondents. Multiple respondents also argued for the use of generation length to assess the gravity of harm to an individual organism when the local population of which it is part is declining. Another example here is the argument that specialisation, in the case of a high local abundance, need not lead to an increase of the gravity score. This can be seen in light of Crisfield et al. (2024), who state that specialisation and rarity do not always go hand in hand. A final example is the disagreement on the use of flagship species as a harm criterion, where some respondents stated that ‘charisma’ does not say anything about functional relevance. However, there are many examples of species that are functionally extinct but do carry great weight in conservation efforts at high financial costs (Elliott et al, 2001). In the end, the importance of the loss of such an individual may perhaps depend on the conservation campaigning context. Ultimately, flagship species are often selected for their ability to capture public attention or as rallying points to stimulate conservation awareness (Caro, 2010). The key aspect of a flagship species’ effectiveness is the extent to which it builds attitudinal, behavioural, financial, or political support and they typically need not have ecological significance as well (Verissimo et al., 2011). The question is whether harming an individual of a flagship species, due to its role in conservation campaigns in a specific region, justifies an increased gravity score in that region (and not elsewhere). If the species also has ecological significance, this should be reflected in other gravity criteria. These examples highlight the importance of adequately contextualizing the discussion of potential gravity criteria. Disagreement about their use may stem from specific caveats rather than their irrelevance. Therefore, several of the criteria that lacked consensus probably deserve further attention.

As such, it is essential for individual criteria to be clear and well-defined, particularly when the information they convey is used in a legal context where decisions based on this information may have far-reaching consequences for offenders. This also implies avoiding overlap of criteria in a decision tool that combines and integrates these criteria (IUCN, 2022; Desair et al., 2024). For example, many respondents viewed generation length and age at first maturation as substitutable harm criteria, or mentioned that population reductions are scaled by the IUCN with generation length to establish the Red List status of species. Similarly, it may be questionable to increase the gravity score twice, based on both the ‘location’ (protected area) and ‘natural integrity’ criteria as sometimes the latter may be the reason an area was granted protection in the first place. This underscores the importance of ensuring that each criterion used in a decision tool provides distinct information, avoiding redundancy or duplication. Ultimately, when criteria overlap, or data is inadequate or ambiguous, this could lead to error propagation and amplification of uncertainty in an overarching gravity score, or to an artificial inflation of the overall gravity of the offence which could in turn provide a basis for excessive sanctioning or imposed over-remediation or -compensation requirements.

Importantly, most respondents highlighted the significance of maintaining ecological baseline conditions as intact as possible, and preferred primary or in-kind remediation with short time lags to limit or partially undo the harm caused by the offence. This most closely aligns with a holistic approach, remediating harm of not just biodiversity’s ecological value but also local communities’ instrumental and relational values assigned to nature that were present prior to the offense (Maron et al., 2016; Moilanen & Kotiaho, 2018; Cole et al., 2021; Tupala et al., 2022). Another reason to why a focus on restoring baseline conditions is called for is that this may afford the prevention of cumulative impacts and shifting baselines (Sala, 2000; Papworth et al, 2009; Soga & Gaston, 2018). Correspondingly, significantly increasing the gravity score for causing irreversible harm to the maximum acceptable levels cannot only guarantee a proportionate compensation for the damage inflicted, but also create a strong deterrent against causing harm in the first place. Moreover, insufficient knowledge of remediation measures and remediation uncertainty are deemed relevant by our respondents to take into account as gravity criteria. The use of gravity criteria to cover for remediation risks and shortcomings is very similar to the use of complementary measures in Environmental Impact Assessment to mitigate or compensate for damage caused by development projects (zu Ermgassen et al., 2019a). For example, similar to such complementary measures, remediation criteria can be used to increase the area of habitat required to mitigate the harm caused or to justify higher financial compensation when in-kind remediation is not possible or carries high risk, such as in areas with a long history of continuous natural integrity (Croshner et al., 2019).

Several respondents noted that while combining various criteria to more accurately assess the gravity of biodiversity offences is desirable, translating these assessments into grades of gravity could quickly become too complex or subjective. The same goes for a differential weighting of criteria. For example, the survey results show how a majority of respondents were of the opinion that aesthetic and cultural values assigned to nature could play a role in the assessment of the gravity of biodiversity offences, but that these should be ‘outweighed’ by ecological and intrinsic concerns. Other stakeholders may see this differently. Therefore, even when an overarching score is considered adequately derived, it is important to clearly distinguish and disclose the different criteria that contributed to it. Additionally, the division into grades of gravity and the weighting of criteria should be transparent. This practice prevents precisely those black-box outcomes that undermine the transparency of results and legitimacy of decisions (Naves et al., 2020; Desair et al., 2024). Here it needs to be said that several respondent also argued that, while more precise and context-specific considerations are important, this may also lead to a lack of standardization, which means more time and expertise (e.g. what criteria to use and why) must be invested in the assessment and in the communication of its outcomes to stakeholders. A solution could be to balance specific and general criteria. Some respondents suggested that this balance is context-dependent and that it is advisable to use a more specific or nuanced analysis in the case the harmed habitat or population is deemed more important for nature conservation or when standardized methods prove inadequate in capturing critical aspects necessary for a comprehensive assessment of the offence gravity. All of the above suggests that, to be effective, the design of a decision tool may need to prioritise practicality and simplicity for operational use. Hence, the question becomes what simplification is acceptable.

Lastly, as is made more clear in the final section, many respondents found it quite challenging to assess the gravity of harm done to an individual organism. This is understandable in light of conservation’s conventional emphasis on preserving ecological collective entities, where individuals are seen primarily as units of a population or species. Indeed, as elaborated on by Marris (2021), comparing the value of individual beings with that of complex ecosystems is in itself a convoluted problem. However, it is important to understand that even within the conventional emphasis, laws protecting individual organisms and sanctioning violations serve as a pragmatic approach to ensure the overall health and viability of populations and species. These laws deter and enable the sanctioning of harmful behaviours like illegal hunting, which, if left unchecked, could cumulatively harm populations (ICCWC, 2022).

### 4.2. Relevance of gravity criteria: implications from value pluralism

The respondents highlighted key issues regarding the valuation of biodiversity at the intersection of legal sciences, environmental ethics and conservation biology which are relevant for determining the gravity of offences. For example, only a minority (11%) of respondents agree the gravity score of an offence should be lowered in case an individual of a species which is functionally extinct is harmed. This could reflect an ethical stance that every individual of a species matters or, even if functionally extinct, a species holds symbolic or intrinsic value. This also aligns with critics who note that the loss of rare species will not always visibly impact ecosystem stability and that a mere instrumental focus would make a portion of biodiversity expendable (Rolston, 1985; Thompson, 2010). Also, nearly half of respondents find it is still justified to increase the gravity score for harming individuals, despite the presence of individuals of other species that have the same roles within the community.

Furthermore, together with the explicit mentioning of nature’s intrinsic value, just 39% of respondents agree that only the population-relative number of harmed individuals should be taken into account. Respondents’ answers also reveal the significance of non-material values of nature such as recreational enjoyment, aesthetic appreciation, educational enrichment, and cultural identity. Additionally, a high majority agreed that when areas with a long history of continuous natural integrity are harmed it is appropriate to increase the gravity score, while pointing at these areas as having intrinsic, historical or symbolic meaning.

Therefore, the findings of our survey indicate the presence of relational and intrinsic arguments to value biodiversity, confirming that there can be multiple, coexisting values assigned to nature (Mace, 2014, Piccolo et al., 2022) as well as that these could influence the individual-based assessment of the gravity of offences. Indeed, not only conservation biologists but also environmental ethicists advocate for reinvigorating the notion of value pluralism and more inclusive legal frameworks, arguing it is our moral responsibility to preserve all species and ecosystems and that this is not just something we are required to do out of narrow self-interest. (Rolston, 1985, Soulé, 1985; Kopnina et al., 2018a; Muradian & Gómez-Baggethun, 2021; Piccolo et al., 2022). Such prescriptive notions can make inroads into reinterpreting the gravity of offences (Garmestani et al., 2019). For example, they may go beyond a prioritization of the material services to human wellbeing nature provides (Washington et al., 2021; Kadykalo et al., 2019) and are increasingly reflected in legal frameworks that shows how intrinsic valuation of nature can ground the moral standing of a natural entity which then in turn is ascribed rights (Bataille et al., 2020; Laastad, 2020; den Outer, 2023).

Viewing biodiversity as merely providing instrumental or economic value may therefore result in a too narrow interpretation of nature’s value and a perception of the gravity of offenses that is too lenient. Many scholars argue it is precisely such anthropocentric reasoning that contributes to the biodiversity crisis and further erodes our connection with nature (Ives & Kendal, 2014; Arias-Arévalo et al., 2018; Jax et al., 2018; Kopnina et al., 2018b; Van den Born et al., 2018; Jacobs et al., 2020; Mattijssen et al., 2020). Conversely, recognizing and incorporating relational and intrinsic values, alongside instrumental values, can lead to a more comprehensive and proportionate judgment of biodiversity offences, reflecting a deeper appreciation of the multifaceted significance of biodiversity (IPBES,2022). This is important, as different underlying reasons to protect nature are easily overlooked despite having the potential of influencing conservation related decision-making and hampering the cooperation between stakeholders (Rowell, 2020; Ghijselinck et al., 2023; Ruales et al., 2024).

Finally, while the practical development of a quantitative decision tool aimed at the individual-based assessment of the gravity of biodiversity offences is undoubtedly important, it is beyond the scope of this paper. Instead, our goal is to provide foundational principles and building blocks necessary to support future work in this area. In this regard, our study can serve as the groundwork for researchers, decision-makers or practitioners who may wish to investigate how harm or remediation criteria can be combined with other, more legal or other gravity criteria. For example, different jurisdictions and political environments have unique legal frameworks, and a decision tool may want to also incorporate recidivism, whether the offence was committed within a professional capacity, intent, or commercial motivation, or prefer to give more weight to certain underlying values of nature compared to others (Wilson & Boratto, 2020, Hutchinson et al., 2023). Moreover, the mode of response to biodiversity offences deserves more investigation. A review from Hutchinson et al. (2023) showed that the predominant response to biodiversity offences is fines, which they argue are ineffective to deter corporations or powerful individuals, nor do such financial penalties anything to repair the damage done. Gravity criteria sustaining an individual-based assessment of biodiversity offences can, by more adequately documenting the harm done and values lost, support the discussion on what effective responses to these offences could be. While not discussed here, the nexus of animal welfare legislation and conservation laws may be interesting to explore where the gravity of offences against individual wild organisms is concerned (Marris, 2021). Integration of animal welfare considerations in international rules relating to biodiversity and conservation is slowly emerging (ICCWC,2022; Katz, 2024). However, as Palmer (2004) and Johnson et al. (2019) show, the degree of gravity of killing individual organisms can be bitterly contested where this pertains to control and eradication of non-native individual organisms which are (considered) harmful to native species. Ultimately, effectively judging the gravity of biodiversity offences demands a synergistic approach bridging different disciplines, not only to ensure an accurate and more well-rounded assessment of ecological harm but also to strengthen the enforcement strategies crucial for safeguarding our planet’s biodiversity heritage.

## Supporting information

Survey

## Notes

### Competing Interest Statement

The authors have declared no competing interest.

