## Supplementary material for "Strengthening conservation law enforcement: incorporating harm, remediation and plural nature value criteria into an individual-based assessment of biodiversity offence gravity": Survey

### Gravity of biodiversity offences

**Thank you for participating in our survey. We appreciate your feedback!**

Your responses will be analysed and used in a study that aims to identify biological, ecological and contextual criteria that adequately capture the gravity (severity) of biodiversity offences and the consequences this has for local, regional and international conservation. With gravity, we refer to the degree of different kinds of harm that biodiversity offences cause.

The intention is to inform the judiciary and administrative authorities and help shape prosecution and sentencing policies, or the sanctioning of biodiversity offences, in a proportional and dissuasive way. Sanctioning refers to the enforcement phase that begins once the law violation has been formally established.

By taking part in the survey you acknowledge that you:
- understand participation is voluntary and you can withdraw at any time without giving a reason
- understand that any data provided cannot be withdrawn once it has been submitted online
- understand that only the researchers will have access to the identities of the participants and the data

**Thank you in advance for your time and effort!**

---

For what type of institute are you currently working in ecology, biology or conservation?

- ☐ University (1)
- ☐ Research institute (2)
- ☐ Government (3)
- ☐ Other (4) ________________________________________________

---

How long have you been involved in ecology, biology or conservation related work or research?

- ◯ 0 - 2 years (1)
- ◯ 2 - 5 years (2)
- ◯ 5 - 10 years (3)
- ◯ > 10 years (4)

---

May we contact you in the future for follow-up questions?

- ◯ No (1)
- ◯ Yes (2)

In the following questions **“gravity score”** is used to refer to the degree of severity of biodiversity loss or harm and serves to create a more unified structure for the survey. A higher gravity score may then be linked to higher penalties or fines, confiscations, and non-custodial approaches such as community service, re-education, remediation measures, or financial compensation.

Often a topic or scenario is introduced by a short explanatory text that is followed by a set of multiple choice questions which have the same primary number.

Most open-ended questions serve to give you the opportunity to elaborate on your answers. While any elaboration provided by you would be highly appreciated, these open-ended questions *(*starting with *please feel free...)* are facultative and leaving them unanswered will not prevent you from ending the survey.

---

1. Consider the loss of **a single Common Fire salamander** (Salamandra salamandra) due to an offence in a Natura 2000 alluvial forest. This species has 3 statuses. It has a **Regional Conservation status** of **Critically Endangered** and a **Global Conservation status** of **Least Concern**. This salamander is also a **Habitat Typical species**, which means its conservation status on a regional level is used to determine whether or not the status of the alluvial forest habitat type is favourable.

---

1.1. Please indicate to what extent you deem the use of the **Regional Conservation status** appropriate in determining the gravity of loss of the salamander.

- Extremely inappropriate (1)
- Somewhat inappropriate (2)
- Neither appropriate nor inappropriate (3)
- Somewhat appropriate (4)
- Extremely appropriate (5)

---

1.2. Please indicate to what extent you deem the use of the **Global Conservation status** appropriate in determining the gravity of loss of the salamander.

- extremely inappropriate (1)
- Somewhat inappropriate (2)
- Neither appropriate nor inappropriate (3)
- Somewhat appropriate (4)
- Extremely appropriate (5)

---

1.3. Please indicate to what extent you agree with the following statements.

*Because it is a **Habitat Typical species** the loss of salamanders may ultimately have a negative impact on the conservation status of the habitat type for which it is typical. This should further increase the gravity score.*

- ◯ Strongly disagree (1)
- ◯ Somewhat disagree (2)
- ◯ Neither agree nor disagree (3)
- ◯ Somewhat agree (4)
- ◯ Strongly agree (5)

---

*1.4. The offence occured in a **Nature Protection Area** (nature reserve, Natura 2000, National Park). This should further increase the gravity score.*

- ◯ Strongly disagree (1)
- ◯ Somewhat disagree (2)
- ◯ Neither agree nor disagree (3)
- ◯ Somewhat agree (4)
- ◯ Strongly agree (5)

---

2. Imagine the loss of an individual of an unspecified, protected species of which **local population trends are now decreasing**. Whatever the regional or global conservation status, to what extent should the gravity score increase compared to when the local population trends are not decreasing?

- not at all (1)
- a little (2)
- somewhat (3)
- to a large extent (4)
- to a great extent (5)
- no opinion (6)

---

3. Imagine the individual lost belongs to a species with favourable local conservation status. Based on status alone the gravity score is therefore low. To what extent should the gravity score increase when the species is threatened **in other parts of its range**?

- not at all (1)
- a little (2)
- somewhat (3)
- to a large extent (4)
- to a great extent (5)
- no opinion (6)

---

4. Please feel free to elaborate on the use of conservation status or local population trends in the determination of a gravity score for an offence.

________________________________________________________________

---

5. **Generation length** is the average time between births of two consecutive generations in a population and may be an indicator of the time needed for the population to recover from the loss of individuals.

Please indicate to what extent you agree with the following statements.

---

*5.1. Taking generation length into account in the determination of a gravity score is important when the individual lost belongs to a local population* ***which is declining****.*

- ○ Strongly disagree (1)
- ○ Somewhat disagree (2)
- ○ Neither agree nor disagree (3)
- ○ Somewhat agree (4)
- ○ Strongly agree (5)

---

*5.2. As an alternative for generation length, the age at which individuals first produce offspring in the wild* ***(age of first reproduction)*** *can be used as a gravity criterion.*

- ○ Strongly disagree (1)
- ○ Somewhat disagree (2)
- ○ Neither agree nor disagree (3)
- ○ Somewhat agree (4)
- ○ Strongly agree (5)

---

5.3. Please feel free to elaborate on the use of generation length to distinguish between the gravity of loss of individuals.

________________________________________________________________

6. Many species are distributed as metapopulations, which are local populations linked by occasional dispersal of individuals or propagules. A **good dispersal ability** is an important species characteristic to maintain viable metapopulations.

Please indicate to what extent you agree with the following statements.

---

*6.1. Poor dispersers are, **in general**, more vulnerable to changes in their habitat, whether this habitat is currently fragmented or not. Therefore, it is justified to increase the gravity score for the loss of an individual of poorly dispersing species.*

- ○ Strongly disagree (1)
- ○ Somewhat disagree (2)
- ○ Neither agree nor disagree (3)
- ○ Somewhat agree (4)
- ○ Strongly agree (5)

---

*6.2. Gravity scores should be increased for activities that **fragment a species' habitat**, regardless of the dispersal ability of the affected species.*

- ○ Strongly disagree (1)
- ○ Somewhat disagree (2)
- ○ Neither agree nor disagree (3)
- ○ Somewhat agree (4)
- ○ Strongly agree (5)

---

*6.3. There should be no increase in gravity score for the loss of a poor disperser when the* ***population density*** *in the affected habitat patch is high.*

- ○ Strongly disagree (1)
- ○ Somewhat disagree (2)
- ○ Neither agree nor disagree (3)
- ○ Somewhat agree (4)
- ○ Strongly agree (5)

---

6.4. Please feel free to elaborate on the use of **dispersal ability** in the determination of a gravity score.

________________________________________________________________

---

7. An important factor to consider in determining a gravity score is **the number** of individuals impacted. This can be an **absolute** number, but also **relative** to the size of the population.

Please indicate to what extent you agree with the following statements.

---

*7.1. Relative numbers should be* ***combined*** *with absolute numbers as only then the value of every individual is counted.*

- Strongly disagree (1)
- Somewhat disagree (2)
- Neither agree nor disagree (3)
- Somewhat agree (4)
- Strongly agree (5)

---

*7.2. Absolute numbers should be given a higher weight than relative numbers.*

- Strongly disagree (1)
- Somewhat disagree (2)
- Neither agree nor disagree (3)
- Somewhat agree (4)
- Strongly agree (5)

---

*7.3. Only the relative number of individuals should be taken into account since this tells us something about the impact on the population which is more important than the persistence of individuals.*

- Strongly disagree (1)
- Somewhat disagree (2)
- Neither agree nor disagree (3)
- Somewhat agree (4)
- Strongly agree (5)

---

7.4. Please feel free to elaborate on the use of relative or absolute numbers of individuals in the determination of a gravity score.

________________________________________________________________

---

8. Consider the following statements about birds. Please indicate to what extent you agree with the following statements.

*Taking into account the loss of reproductive fitness, the gravity score should be higher for the loss of a parent bird than for a fledgling.*

- Strongly disagree (1)
- Somewhat disagree (2)
- Neither agree nor disagree (3)
- Somewhat agree (4)
- Strongly agree (5)

---

8.1. Please feel free to elaborate on this statement.

________________________________________________________________

*8.2. Regardless of reproductive status, the gravity score should be higher for a mature than for an immature bird.*

- ○ Strongly disagree (1)
- ○ Somewhat disagree (2)
- ○ Neither agree nor disagree (3)
- ○ Somewhat agree (4)
- ○ Strongly agree (5)

8.3. Please feel free to elaborate on this statement.

________________________________________________________________

*8.4. The gravity score for the loss of a parent bird should be higher in case of uniparental care than in the case of biparental care.*

- ○ Strongly disagree (1)
- ○ Somewhat disagree (2)
- ○ Neither agree nor disagree (3)
- ○ Somewhat agree (4)
- ○ Strongly agree (5)

8.5. Please feel free to elaborate on this statement.

________________________________________________________________

*8.6. The loss of a fledgling or a parent should lead to a higher gravity score in a single-brooded versus a multi-brooded species.*

- ○ Strongly disagree (1)
- ○ Somewhat disagree (2)
- ○ Neither agree nor disagree (3)
- ○ Somewhat agree (4)
- ○ Strongly agree (5)

8.7. Please feel free to elaborate on this statement.

________________________________________________________________

9. **Natural dune landscapes** contain the following dune habitats:

**1.** Dynamic dunescapes with shifting sands.

**2.** Wet dune slacks, which occur as small, naturally fragmented systems in the dune landscape. These are of great importance to many specialist fauna and flora species such as Marsh grass of Parnassus (Parnassia palustris) or Musk Orchid (Herminium monorchis) which often find little or no suitable habitat outside these areas.

**3.** Fixated dunescapes such as dune scrub with sea-buckthorn (Hippophae rhamnoides) and wooded dunes.

Most of these dune habitats are now designated Natura 2000-sites but have become **small, isolated and enclosed in an urbanised landscape in some regions**. Protected zones are therefore interspersed with some others with no protection. Moreover, it has been established that habitat fragmentation stimulates fixation of dune slacks and leads to lower fitness and reduced genetic diversity of the remaining specialised plant populations.

Now, imagine a case of illegal construction or land use in one of the less protected zones. Please indicate to what extent you agree with the following statements.

---

*9.1. The loss of **habitat specialists** should be given more weight when determining a gravity score when compared to the one given to **generalists**.*

- ○ Strongly disagree (1)
- ○ Somewhat disagree (2)
- ○ Neither agree nor disagree (3)
- ○ Somewhat agree (4)
- ○ Strongly agree (5)

---

9.2. Please feel free to elaborate on this statement.

________________________________________________________________

---

*9.3. Potential **indirect impacts** from illegal land use in the unprotected site (related to fragmentation, dune fixation) **on habitat specialists** in the Natura 2000 dune slacks should be taken into account when determining a gravity score.*

- ○ Strongly disagree (1)
- ○ Somewhat disagree (2)
- ○ Neither agree nor disagree (3)
- ○ Somewhat agree (4)
- ○ Strongly agree (5)

---

9.4. Please feel free to elaborate on this statement.

________________________________________________________________

*9.5. When illegal activities lead to* ***reduced dune dynamics*** *and* ***accelerated fixation*** *the gravity score of impacts of these activities should be increased.*

- ◯ Strongly disagree (1)
- ◯ Somewhat disagree (2)
- ◯ Neither agree nor disagree (3)
- ◯ Somewhat agree (4)
- ◯ Strongly agree (5)

*9.6. Compensation measures for illegal land use should account for the negative impact on the* ***loss of connectivity*** *between protected habitat patches or increased isolation of protected areas,* ***even when these happen in unprotected zones****.*

- ◯ Strongly disagree (1)
- ◯ Somewhat disagree (2)
- ◯ Neither agree nor disagree (3)
- ◯ Somewhat agree (4)
- ◯ Strongly agree (5)

9.7. Please feel free to elaborate on this statement.

________________________________________________________________

*9.8. A **limited dispersal capacity** of the habitat specialists impacted (such as plant species in dune slacks) should be seen as an **additional gravity factor** (on top of specialism).*

○ Strongly disagree (1)

○ Somewhat disagree (2)

○ Neither agree nor disagree (3)

○ Somewhat agree (4)

○ Strongly agree (5)

---

9.9. Please feel free to elaborate on this statement.

________________________________________________________________

---

10. Take into consideration the following concepts:

**Keystone species:** a species whose influence(s) on ecological processes is very important, and greater than one would expect on the basis of its abundance or biomass alone.

**Umbrella species:** a species whose conservation indirectly protects many other species in the ecosystem and is used to make conservation-related decisions.

**Naturally rare species:** a species at risk which is characterized by small or isolated populations as a result of its limited geographic range, small ecological niche, and/or low reproductive rate.

**Artificially rare species:** a species whose abundance and/or distribution has been strongly reduced and is only rare due to anthropogenic disturbance.

**Functionally extinct species:** a species that no longer plays any significant role in ecosystem function.

Please indicate to what extent you agree with the following statements.

---

*10.1. Harming an individual belonging to a* ***keystone species*** *should lead to a higher gravity score compared to other species.*

- ○ Strongly disagree (1)
- ○ Somewhat disagree (2)
- ○ Neither agree nor disagree (3)
- ○ Somewhat agree (4)
- ○ Strongly agree (5)

---

10.2. Please feel free to elaborate on this statement.

________________________________________________________________

*10.3. Harming an individual belonging to an* ***umbrella species*** *should receive a higher gravity score compared to other species.*

- Strongly disagree (1)
- Somewhat disagree (2)
- Neither agree nor disagree (3)
- Somewhat agree (4)
- Strongly agree (5)

10.4. Please feel free to elaborate on this statement.

- Click to write Choice 1 (1)
- Click to write Choice 2 (2)
- Click to write Choice 3 (3)

*10.5. The gravity score should be higher for harming an individual from an* ***artificially rare species*** *compared to a* ***naturally rare species****, because the former is not evolutionarily adapted to rarity.*

- Strongly disagree (1)
- Somewhat disagree (2)
- Neither agree nor disagree (3)
- Somewhat agree (4)
- Strongly agree (5)

10.6. Please feel free to elaborate on this statement.

________________________________________________________________

*10.7. The gravity score for the loss of individuals should be lower when individuals from another species that fulfil similar roles in the ecosystem remain present.*

- ○ Strongly disagree (1)
- ○ Somewhat disagree (2)
- ○ Neither agree nor disagree (3)
- ○ Somewhat agree (4)
- ○ Strongly agree (5)

10.8. Please feel free to elaborate on this statement.

________________________________________________________________

*10.9. When the loss of individuals concerns a species which is* ***functionally extinct*** *the gravity score should be lower.*

- ○ Strongly disagree (1)
- ○ Somewhat disagree (2)
- ○ Neither agree nor disagree (3)
- ○ Somewhat agree (4)
- ○ Strongly agree (5)

10.10. Please feel free to elaborate on this statement.

________________________________________________________________

11. Consider the use of species richness (the number of species in an area or habitat patch) as a metric to assess the gravity score of habitat loss. Please indicate to what extent you agree with the following statements.

11.1. *To assess harm done to a habitat, indicators of changes in* ***species richness*** *can be used.*

- ○ Strongly disagree (1)
- ○ Somewhat disagree (2)
- ○ Neither agree nor disagree (3)
- ○ Somewhat agree (4)
- ○ Strongly agree (5)

11.2. Please feel free to elaborate on the use of species richness as a criterion to assess the gravity score of harming habitats.

________________________________________________________________

12. The gravity of habitat loss may be, beyond its size, affected by characteristics related to **habitat quality**, or its **refuge function**.

Please indicate to what extent you agree with the following statements.

*12.1. The gravity score of forest loss should be increased according to the presence of **ancient forest plant species**.*

- ○ Strongly disagree (1)
- ○ Somewhat disagree (2)
- ○ Neither agree nor disagree (3)
- ○ Somewhat agree (4)
- ○ Strongly agree (5)

---

12.2. Please feel free to elaborate on this statement.

________________________________________________________________

---

*12.3. When a lost ancient forest area is replaced by planting a **forest of the same size**, the lost ecological value of the mature forest is compensated for.*

- ○ Strongly disagree (1)
- ○ Somewhat disagree (2)
- ○ Neither agree nor disagree (3)
- ○ Somewhat agree (4)
- ○ Strongly agree (5)

---

12.4. Please feel free to elaborate on this statement.

________________________________________________________________

---

*12.5. The more time needed for a replanted forest to reach an equivalent ecological state of the forest that is lost, the more the gravity score should increase.*

- ◯ Strongly disagree (1)
- ◯ Somewhat disagree (2)
- ◯ Neither agree nor disagree (3)
- ◯ Somewhat agree (4)
- ◯ Strongly agree (5)

12.6. Please feel free to elaborate on this statement.

________________________________________________________________

*12.7. Consider the loss of an unspecified native vegetation. When this is* ***permanent*** *- in that it cannot be restored - this should increase the gravity score even when this removal happens in zones that have no protected status.*

- ◯ Strongly disagree (1)
- ◯ Somewhat disagree (2)
- ◯ Neither agree nor disagree (3)
- ◯ Somewhat agree (4)
- ◯ Strongly agree (5)

12.8. Please feel free to elaborate on this statement.

________________________________________________________________

*12.9. The gravity score should be increased when an area or site with a* ***long history of continuous natural integrity*** *is impacted (e.g., old-growth forest, freely running river system).*

- ◯ Strongly disagree (1)
- ◯ Somewhat disagree (2)
- ◯ Neither agree nor disagree (3)
- ◯ Somewhat agree (4)
- ◯ Strongly agree (5)

---

12.10 Please feel free to elaborate on this statement.

________________________________________________________________

---

*12.11. When the loss concerns nature elements such as hedgerows, trees or shrubs* ***that may facilitate dispersal*** *between patches of habitat, the gravity score should be increased beyond that obtained through the inclusion of other factors.*

- ◯ Strongly disagree (1)
- ◯ Somewhat disagree (2)
- ◯ Neither agree nor disagree (3)
- ◯ Somewhat agree (4)
- ◯ Strongly agree (5)

---

13. Concepts such as **remediation** or **ecological equivalence** lie at the heart of compensating for biodiversity losses but the **location of remediation measures**, **interim losses**, **uncertainty** and the **frame of reference** against which this is to be achieved remain important methodological issues.

Please indicate to what extent you think it is appropriate **to increase the gravity score** in the following cases.

---

*13.1. When loss or harm is hard to compensate for* ***in-kind,*** *which means that any individual entities lost or harmed cannot be compensated for through replacement with the same entities (species, vegetation, habitat types, biotopes etc).*

- ◯ Extremely inappropriate (1)
- ◯ Somewhat inappropriate (2)
- ◯ Neither appropriate nor inappropriate (3)
- ◯ Somewhat appropriate (4)
- ◯ Extremely appropriate (5)

---

13.2. Please feel free to elaborate on this statement.

________________________________________________________________

---

*13.3. When a natural zone is lost and is remediated by restoring or protecting habitats* ***elsewhere (compared to locally),*** *because local people lose the benefits this nearby natural zone provided.*

- ○ Extremely inappropriate (1)
- ○ Somewhat inappropriate (2)
- ○ Neither appropriate nor inappropriate (3)
- ○ Somewhat appropriate (4)
- ○ Extremely appropriate (5)

13.4. Please feel free to elaborate on this statement.

________________________________________________________________

*13.5. When harming biodiversity cannot be remediated within a reasonable time-frame after the impact.*

- ○ Extremely inappropriate (1)
- ○ Somewhat inappropriate (2)
- ○ Neither appropriate nor inappropriate (3)
- ○ Somewhat appropriate (4)
- ○ Extremely appropriate (5)

13.6. Please feel free to elaborate on this statement.

________________________________________________________________

*13.7. When* ***knowledge of the affected habitats*** *or species is considered insufficient to achieve adequate remediation.*

- ○ Extremely inappropriate (1)
- ○ Somewhat inappropriate (2)
- ○ Neither appropriate nor inappropriate (3)
- ○ Somewhat appropriate (4)
- ○ Extremely appropriate (5)

13.8. Please feel free to elaborate on this statement.

________________________________________________________________

*13.9. When the outcomes of remediation measures for the persistence of valuable local populations/habitat patches are uncertain.*

- ○ Extremely inappropriate (1)
- ○ Somewhat inappropriate (2)
- ○ Neither appropriate nor inappropriate (3)
- ○ Somewhat appropriate (4)
- ○ Extremely appropriate (5)

13.10. Please feel free to elaborate on this statement.

________________________________________________________________

14. The following questions enquire about whether different values assigned to nature are, in your personal opinion, grounds for increasing or decreasing the gravity score of a biodiversity offence.

*14.1. Increasing the gravity score is justified when landscape elements that are* ***locally or regionally recognised as part of natural or cultural heritage*** *are harmed or lost (such as hedgerows, shrubs and historical grasslands that have been part of the agricultural landscape for a very long time).*

- Strongly disagree (1)
- Somewhat disagree (2)
- Neither agree nor disagree (3)
- Somewhat agree (4)
- Strongly agree (5)

14.2. Please feel free to elaborate on this statement.

________________________________________________________________

*14.3. Increasing the gravity score is justified when an individual that belongs to a* ***flagship species*** *- which is a species that is compelling or charismatic to the public - is harmed.*

- Strongly disagree (1)
- Somewhat disagree (2)
- Neither agree nor disagree (3)
- Somewhat agree (4)
- Strongly agree (5)

14.4. Please feel free to elaborate on this statement.

________________________________________________________________

*14.5. Increasing the gravity score is justified when **aesthetic properties** of landscape elements, or **feelings of reverence or awe**, are involved (such as aesthetically appealing and very old, landmark trees).*

- ○ Strongly disagree (1)
- ○ Somewhat disagree (2)
- ○ Neither agree nor disagree (3)
- ○ Somewhat agree (4)
- ○ Strongly agree (5)

14.6. Please feel free to elaborate on this statement.

________________________________________________________________

*14.7. Take the following instrumental values of nature, here presented as human benefits derived from nature:*
***- provisioning services**, such as food, water, energy, medicinal resources*
***- regulating services**, that influence the conditions in which we live, such as climate regulation and water purification, CO2-sequestration, pollination and biological control.*

***Decreasing the gravity score** is justified when harm or loss can be compensated for in such a way that any provisioning and regulating ecosystem services present are upheld, **even when this is not in-kind**.*

- ◯ Strongly disagree (1)
- ◯ Somewhat disagree (2)
- ◯ Neither agree nor disagree (3)
- ◯ Somewhat agree (4)
- ◯ Strongly agree (5)

---

14.8. Please feel free to elaborate on this statement.

________________________________________________________________

---

15. Should the assessment of the gravity of biodiversity offences be based on solely ecological concerns or can social aspects, such as who benefits from compensation and how, be equally important? Please elaborate on your answer.

________________________________________________________________

---

16. It can be argued that a high precision or specificity in assessing the impacts on individual entities or habitats is desirable but that a balance should be struck between the use of specific criteria versus general ones in assessing the gravity of biodiversity offences. What is your opinion on this and do you have suggestions for such metrics?

________________________________________________________________

17. Comprehensive assessments of the gravity of biodiversity offences are intended to reflect the value we assign to nature. However, it can be argued that justifications for this value are often vague and ambiguous. Can you tell us why the conservation of nature is valuable to you personally?
